# The consequences of mixed-species malaria parasite co-infections in mice and mosquitoes for disease severity, parasite fitness, and transmission success

**DOI:** 10.1101/761791

**Authors:** Jianxia Tang, Thomas J. Templeton, Jun Cao, Richard Culleton

## Abstract

The distributions of human malaria parasite species overlap in most malarious regions of the world, and co-infections involving two or more malaria parasite species are common. Little is known about the consequences of interactions between species during co-infection for disease severity and parasite transmission success. Anti-malarial interventions can have disproportionate effects on malaria parasite species, and may locally differentially reduce the number of species in circulation. Thus, it is important to have a clearer understanding of how the interactions between species affect disease and transmission dynamics. Controlled competition experiments using human malaria parasites are impossible, and thus we assessed the consequences of mixed-species infections on parasite fitness, disease severity, and transmission success using the rodent malaria parasite species *Plasmodium chabaudi*, *P. yoelii yoelii*, and *P. vinckei lentum*. We compared the fitness of individual species within single species and co-infections in mice. We also assessed the disease severity of single versus mixed infections in mice by measuring mortality rates, anaemia, and weight loss. Finally, we compared the transmission success of parasites in single or mixed species infections by quantifying oocyst development in *Anopheles stephensi* mosquitoes. We found that co-infections of *P. yoelii* with either *P. vinckei* or *P. chabaudi* led to a dramatic increase in infection virulence, with 100% mortality observed in mixed species infections, compared to no mortality for *P. yoelii* and *P. vinckei* single infections, and 40% mortality for *P. chabaudi* single infections. The increased mortality in the mixed infections was associated with an inability to clear parasitaemia, with the non-*P. yoelii* parasite species persisting at higher parasite densities than in single infections. *P. yoelii* growth was suppressed in all mixed infections compared to single infections. Transmissibility of *P. vinckei* and *P. chabaudi* to mosquitoes was also reduced in the presence of *P. yoelii* in co-infections compared to single infections. The increased virulence of co-infections containing *P. yoelii* (reticulocyte restricted) and *P. chabaudi* or *P. vinckei* (predominantly normocyte restricted) may be due to parasite cell tropism and/or immune modulation of the host. We explain the reduction in transmission success of species in co-infections in terms of inter-species gamete incompatibility.

## Introduction

Eight malaria parasite species are infectious to humans; namely, *Plasmodium falciparum*, *Plasmodium vivax*, *Plasmodium malariae*, *Plasmodium ovale wallikeri*, *Plasmodium ovale curtisii*, *Plasmodium knowlesi*, *Plasmodium cynomolgi,* and *Plasmodium simium*. The latter three species are parasites of non-human primates, but also cause zoonotic malaria in humans [1–3]. In large parts of the tropical world the ranges of at least some of these species overlap, they are often vectored by the same mosquitoes [4], and mixed-species infections are common [5–7].

Mixed species infections of human malaria parasites are well-documented in natural [8–12] and experimental (e.g. [13, 14]) settings. They are studied regarding diagnosis [15–17], treatment [18], immune response [19], virulence [12, 20], transmission [21–23], and in discussions of public health policy [6]. The virulence of malaria infection is also of interest in the context of co-infection with other pathogens, such as HIV and *Schistosoma* [24–26].

The consequences of mixed-species infections on malaria disease and parasite fitness are incompletely understood. There is conflicting evidence from laboratory and field studies regarding the capacity of mixed-species infections to exacerbate [27] or ameliorate [14] disease. Furthermore, the mechanisms underlying the interactions between parasite species in mixed infections are complicated and multi-factorial, possibly involving both within-host competition [28] and cross-immunity [29].

Mixed species and mixed strain *Plasmodium* infections have been studied in primate [30] and rodent malaria parasite models [27, 28, 31, 32], as these enable the study of all parasite lifecycle stages including those that occur in mosquitoes. The rodent model is enhanced by the availability of multiple rodent malaria parasite species; namely, *Plasmodium yoelii*, *Plasmodium berghei*, *Plasmodium vinckei*, and *Plasmodium chabaudi*. For *P. yoelii* there are additionally several parasite strains that differ in virulence following inoculation of mice [33].

Here we describe the results of a series of experiments utilising multiple strains of the rodent malaria parasite species *P. yoelii*, *P. chabaudi*, and *P. vinckei*, in which mixed infections of various combinations of species and strains were established and studied in both mice and mosquitoes. The consequences of mixed species infections for disease severity in both hosts, parasite fitness, and transmission capacity were analysed.

## Materials and methods

### Ethics statement

Animal experimentation was performed in strict accordance with the Japanese Humane Treatment and Management of Animals Law (law No. 105, 1973; modified 2006), and according to the Regulations on Animal Experimentation at Nagasaki University, Japan. All procedures were approved by the Institutional Animal Research Committee of Nagasaki University.

### Parasites, mice, and mosquitoes

Four rodent malaria parasite strains, comprising three species, were used in these experiments; specifically, *P. chabaudi chabaudi* clone AJ, *P. chabaudi chabaudi* clone AS_ED_ (intermediate virulence, normocyte preference), *P. yoelii yoelii* clone CU (non-virulent, reticulocyte restricted), and *P. vinckei lentum* clone DS (non-virulent, normocyte preference). Six-week-old female CBA mice, *Mus musculus*, were purchased from SLC Inc. (Shizuoka, Japan) and were used for all experiments. Mice were housed in a 12-hour/12-hour light/dark cycle at °C and fed with 0.05% para-aminobenzoic acid (PABA)-supplemented water to assist the growth of parasites. *Anopheles stephensi* mosquitoes were housed in a temperature- and humidity-controlled insectary at 23°C and 75% humidity. Mosquitoes used in the transmission experiments were maintained on 10% glucose solution supplemented with 0.05% PABA.

### DNA extraction and real time quantitative PCR (qPCR)

To determine the proportion of each species in mixed infections, quantitative real time PCR (qPCR) was used to measure copy numbers of the merozoite surface protein 1 gene (*msp1*). DNA was extracted from infected mouse tail blood and infected mosquito midguts using an EZI DNA Investigator Kit (GIAGQN) according to the manufacturer’s instructions. Quantitative PCR was performed on an ABI 7500 real-time PCR machine using a Power SYBR Green kit (Applied Biosystems, UK). Primers were designed based on a species-specific region of *msp1*, as follows: *Py*CUmsp1F 5’CACCCTCAATAAACCCTGC-3’, *Py*CUmsp1R 5’-CGTGTACCAATACTTGAGTCAGAAC-3’; *Pv*DSmsp1F 5’-CAAGAAGCCTCACAACAAGAATCTA-3’, *Pv*DSmsp1R, 5’TGCTGGTTGGGCAGGTGCTGGA-3’ and *Pc*AJmsp1F 5’-GTACAAGAAGGAGCATCAGC-3’, *Pc*AJmsp1R 5’-GCGGGTTCTGTTGAGGCTCCT-3’. PCR assays were conducted on an AB7500 real-time PCR machine (Applied Biosystems, Japan) under the conditions: initial denaturation step of 50°C for 2 min, 95°C for 10 min, followed by 40 cycles of 95°C for 15 sec and finally 61°C for 1 min. Copy numbers of *msp1* were quantified with reference to a standard curve generated from known numbers of plasmids containing the target sequence. As different parasite species have differing mean copy numbers of *msp1* per infected erythrocyte (due to different rates of DNA replication, differing numbers of merozoites per schizont, and differing propensities for multiple erythrocyte invasion), we normalized the proportion of each species in mixed infections by the copy numbers per infected erythrocyte calculated from single species infections. This methodology was also used to quantify the numbers of species-specific oocysts on the midguts of co-infected mosquitoes.

### Experiments involving the monitoring of virulence in mice

Eight experimental groups of five mice each were set up to measure the effects of co-infections of parasite species on mice and to compare the growth of species in single and mixed species infections. Three of these groups were inoculated via intravenous (IV) injection with parasite infected red blood cells (iRBCs) of a single *P. c. chabaudi* clone AJ (hereafter referred to as *Pc*AJ), *P. c. chabaudi* AS_ED_ (*Pc*AS_ED_), *P. y. yoelii* clone CU (*Py*CU) or *P. v. lentum* clone DS (*Pv*DS). The remaining four received mixtures of two species in equal numbers (*Pc*AJ+*Py*CU, *Py*CU+*Pc*AS_ED_*, Py*CU+*Pv*DS and *Pc*AJ+*Pv*DS). Inocula were diluted in a solution of 50% foetal calf Serum (FCS) and 50% Ringer’s solution (27 mM KCl, 27 mM CaCl_2_, 0.15 M NaCl). Mice infected with single parasite species received 10^6^ iRBCs, and co-infected mice received 10^6^ of each component parasite species. It has been shown that a 2-fold difference in parasite numbers has a negligible effect on parasite dynamics and virulence [34].

Virulence was determined using the parameters of mortality, weight loss, and reduction in erythrocyte density, and was measured daily up to day 30 post-inoculation. Erythrocyte densities were counted using a Coulter Counter (Backman coulter, Florida) from 1:40,000 dilution of 2 μl whole blood sampled from tails in Isoton solution (Beckman coulter, Florida). Giemsa’s solution stained thin blood smears from tail vein blood were monitored for parasitaemia for 30 days post-inoculation to assess the parasite replication rate. Whole blood samples (10 μl) were collected daily from day 1 to day 30 into citrate saline, centrifuged briefly, and the erythrocyte pellet stored at −80°C prior to DNA extraction using an EZ1 DNA Investigator Kit (QIAGEN, Japan) and an EZ1 BioRobot (QIAGEN, Japan). Species specific qPCR based on the *msp1* gene was used to measure the proportions of each parasite in the mixed infections [32, 35]. All experiments were performed twice.

### Mosquito transmission experiments: estimation of mosquito fitness and parasite species transmission capacity

Groups of mice were infected with single and mixed species infections of *Py*CU, *Pv*DS, and *Pc*AJ parasites (total 6 groups, each of 5 mice). On days 3 and 5 post-inoculation, individual groups of mosquitoes (n = 40 mosquitoes per group; 5-7 days post emergence from pupae) were fed on individual mice. Immediately following the feed, 20 mosquitoes from each group were pooled by mouse group into 12 cages, and egg bowls added two days later to allow the collection of eggs. These groups were monitored for longevity by counting dead mosquitoes daily up to day 60, and the numbers of larvae produced per mosquito were counted at day 5 post-hatching. Seven days later, 20 mosquitoes were removed from each group (57 groups total, as the number of mice fed from group *Pv*DS and *Pc*AJ+*Pv*DS was reduced to three and four, respectively for the day 5 feed) and the midgut oocyst burden recorded following dissection. Dissected midguts were stored at −80°C prior to DNA extraction for species proportion analysis by qPCR.

### Statistical analyses

All graphs were generated using GraphPad Prism 6 (GraphPad software Inc, USA). Comparison of survival curves was carried out using Log-rank (Mantel-Cox) tests. Multiple t-tests, corrected for multiple comparisons using the Holm-Sidak method, were used for comparing parasitaemia, erythrocyte density, weight loss, and parasite density of single and mixed infection in mice at all days during infection. Mann Whitney tests were carried out for cumulative parasite density, mosquito infection, and analysis of oocysts per gravid mosquito. P-values of below 0.05 were considered significant.

## Results

### Mixed species parasite infections involving a reticulocyte specialist and a normocyte specialist are more virulent and cause greater host mortality than single species infections in mice

Infection parameters for single and mixed infections involving *Py*CU (reticulocyte restricted) and either *Pv*DS, *Pc*AJ, or *Pc*AS_ED_ (normocyte preference) are summarized in Table 1. *Py*CU and *Pv*DS are not lethal in single species infections and only the intermediately virulent species *P. chabaudi* (*Pc*AJ and *Pc*AS_ED_) caused death of mice in single infections, with 40% mortality occurring between days 9 and 13 post-infection (PI) for both *Pc* strains. In contrast, mixed-species co-infections of *Py*CU with either *Pv*DS, *Pc*AJ, or *Pc*AS_ED_ resulted in highly virulent infections with 100% mortality (Fig. 1A-C).

**Fig. 1.**
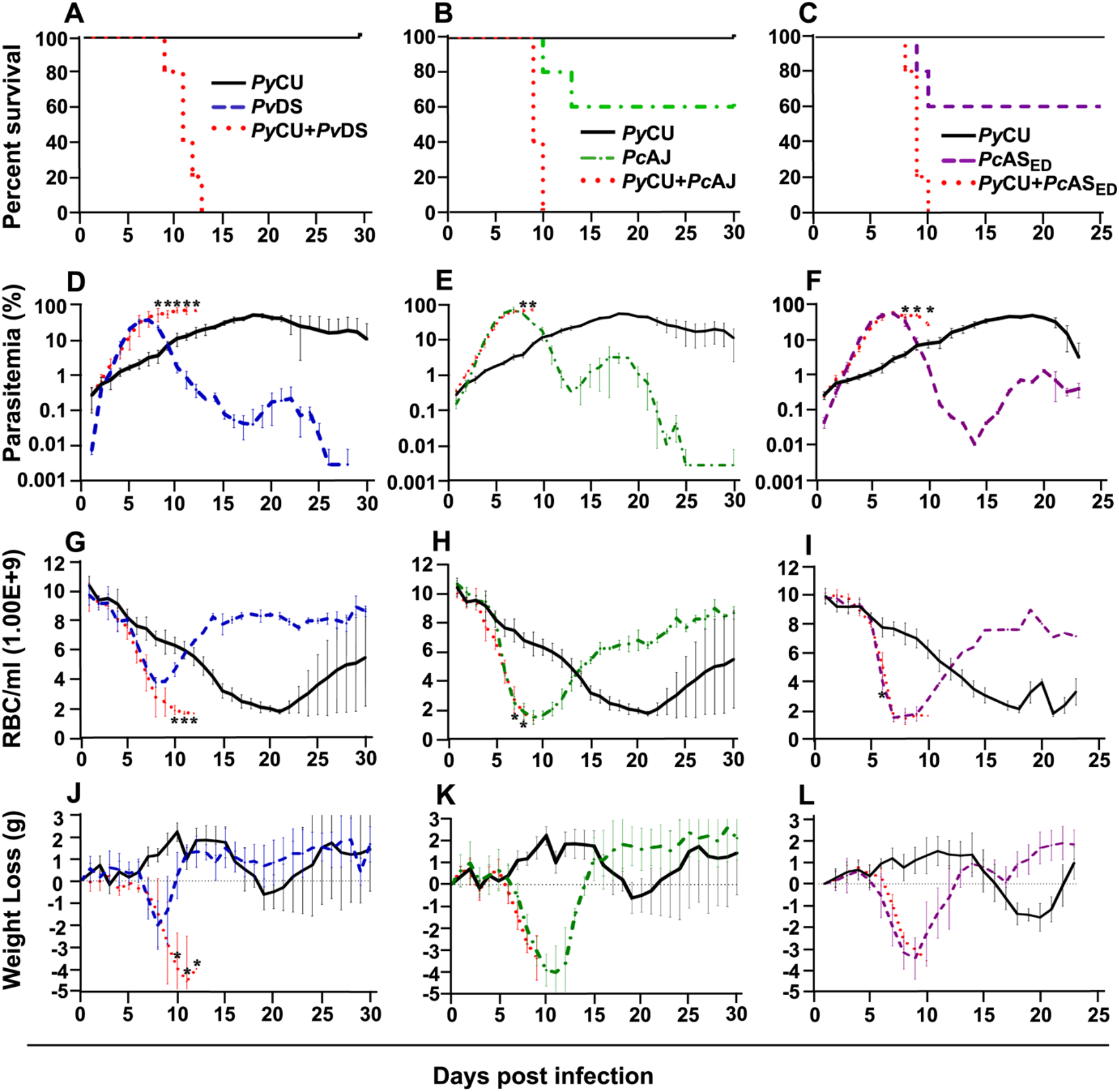
Percent survival (panels A-C), parasitaemia (panels D-F), erythrocyte density (panels G-I), and weight loss (panels J-L) of mice infected with *Plasmodium yoelii* CU (black), *Plasmodium vinckei* DS (blue), *Plasmodium chabaudi* AJ (green), and *Plasmodium chabaudi* AS_ED_ (purple) species in single and mixed infections (red). Mice were inoculated via intravenous injection of 1×10^6^ infected red blood cells of either single or mixed species of the above parasites and followed for 25 to 30 days to determine mortality, live-body weight, erythrocyte density, and parasitaemia. Data points indicate the mean value for five mice in each experimental group and error bars indicate the standard error of the mean (SEM). An asterisk represents statistically significant differences between mixed infections compared with both single infections (Multiple t-tests, with the assumption that all rows are sampled from populations with the same scatter and corrected for multiple comparisons using the Holm-Sidak method). Parasitaemias of *Py*CU and *Pv*DS in mixed-infections were significantly different between days 8 and 12 post infection (PI) compared with the same strains in single infections (panel D); *Py*CU and *Pc*AJ mixed-infections were significantly different to single infections of the same strains on days 8 and day 9 PI (panel E); and *Py*CU and *Pc*AS_ED_ mixed-infections were significantly different from single infections between days 8 and 10 PI (panel F). Erythrocyte densities of *Py*CU and *Pv*DS mixed-infections were significantly different from single infections on day 10 to day 12 PI (panel G); *Py*CU and *Pc*AJ mixed-infections were significantly different on days 7 and 8 PI (panel H); while mixed infections of *Py*CU and *Pc*AS_ED_ were significantly different from single infections only on day 6 PI (panel I). Mice infected with *Py*CU+*Pv*DS mixed-infections lost significantly more weight compared to single infections from day 10 to day 12 (panel J). Mice in the groups infected with mixed infections of *Py*CU+*Pc*AJ suffered from significantly reduced erythrocyte density compared to mice in single infection groups (Two-way RM ANOVA measured mixed effects model, P=0.043, F=5.7, DFn=1, DFd=8). Detailed statistical values relating to significance are given in Supporting Table 1. Experiments were repeated twice, data is from one representative experiment.

The mixed species co-infections were all characterised by protracted parasitaemia and prolonged chronicity of disease compared to single infections of their constituent species. Mixed species infections resulted in higher parasitaemia than either of their constituent species in single infections (Fig. 1D) and peak parasitaemia occurred on the same day PI as the more virulent of the constituent species; except for *Py*CU + *Pv*DS in which peak parasitaemia occurred between days 8-11, compared to the *Pv*DS single infection, in which peak parasitaemia occurred at days 6-7 (Fig. 1D-F). Host mortality in mixed species infections occurred at peak parasitaemia and was presumably caused by anaemia resulting from an inability to clear parasites from the blood.

During the latter stages of the infection mixed species infections involving *Pv*DS and *Py*CU resulted in lower erythrocyte densities (Fig.1 G-I) and greater weight loss (Fig.1 J-L) compared to either of the constituent species in single infections. This increased pathology was linked to the inability of mice to control parasitaemia.

### Mixed species parasite infections involving two normocyte specialists result in protracted parasitaemia, but not increased virulence

*P. vinckei* DS and *P. chabaudi* AJ are both normocyte invading parasites. *Pc*AJ is of moderate virulence, causing rapid and severe anaemia and weight loss during the first 10 days of infection when parasitaemia rises, and results in 40% host mortality (Table 2 and Fig. 2A). *Pv*DS is a much less virulent parasite, causing less severe weight loss and milder anaemia, and is never lethal (Table 2 Fig. 2A). The combination of these two parasites in a mixed infection results in 50% host mortality, and a pathology consistent with that of *Pc*AJ, the more virulent of the two species (Table 2 and Fig. 2A). However, the *Pc*AJ+*Pv*DS infection results in three distinct parasitaemia peaks, compared to the two peaks produced by the single species, and parasitaemias persisted up to the last day of the experiment (day 30), compared to clearance by day 26 in the single species infection (Fig. 2B). The *Pc*AJ+*Pv*DS infection also displayed a sharper decline in parasitaemia following the first peak (Fig. 2B), and this was associated with a quicker recovery from anaemia and weight loss between days 10 and 15 (Fig. 2C, D).

**Fig. 2.**
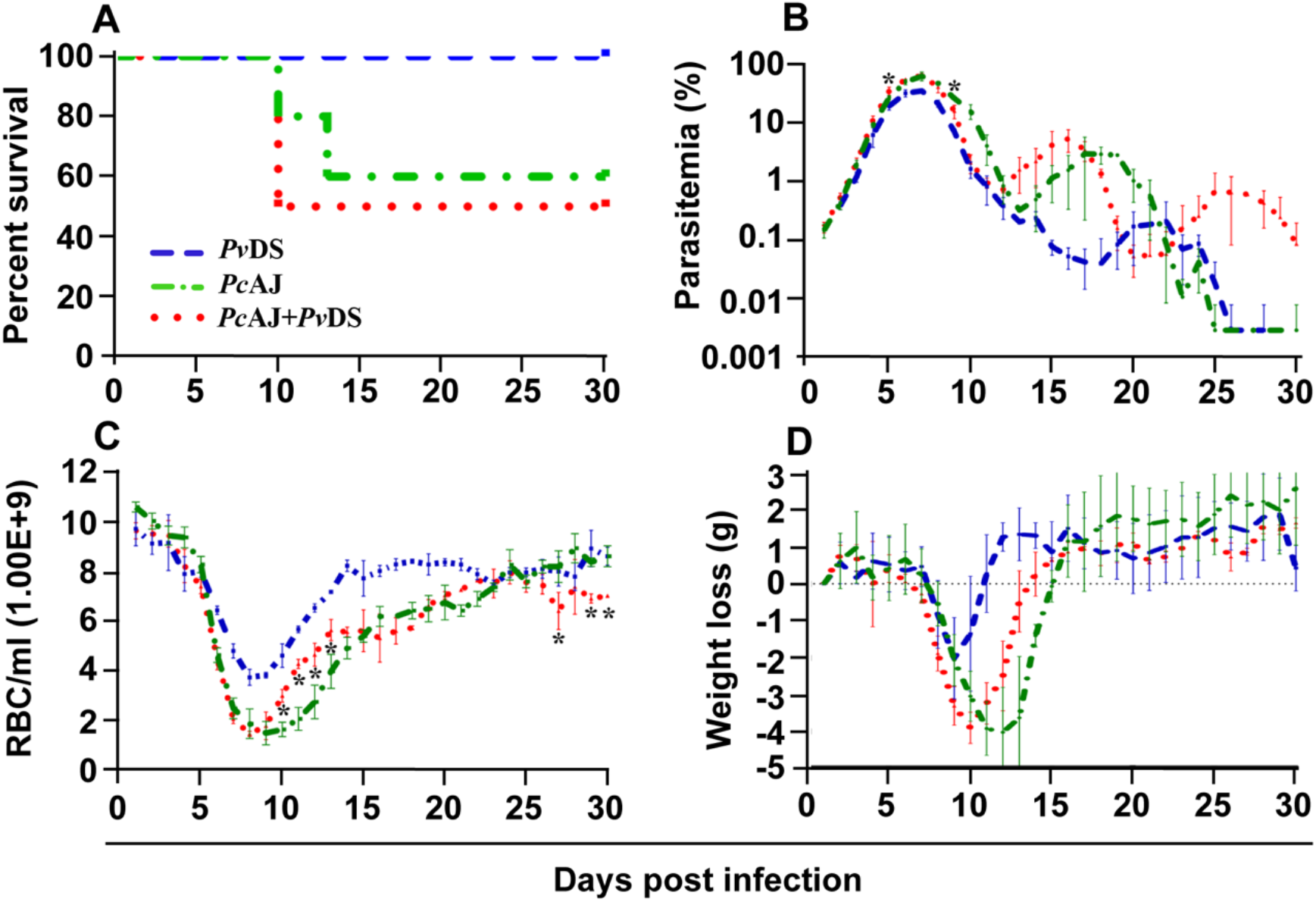
Percentage survival (panel A), parasitaemia (panel B), erythrocyte density (panel C), and weight loss (panel D) of infected mice with *Plasmodium vinckei* DS (blue), *Plasmodium chabaudi* AJ (green) single or mixed infections (red). Mice were inoculated via intravenous injection of 1×10^6^ infected red blood cells of either single or mixed species of the above parasites and followed for 30 days to determine mortality, live-body weight, erythrocyte density, and parasitaemia. Data points indicate the mean value for mice of each experimental group and error bars indicate the standard error of the mean (SEM). An asterisk represents statistically significant difference of mixed infections compared with both single infections (Multiple t-tests, with the assumption that all rows are sampled from populations with the same scatter and corrected for multiple comparisons using the Holm-Sidak method). Mice infected with mixed-infections developed statistically significantly higher parasitaemia on days 5 and 9 PI compared with both single infections (panel B). Mice infected with mixed-infections had significantly lower erythrocyte densities on days 10 to day 13, day 27, day 29 and day 30 PI compared with both single infections (panel C). Considering the entire time course of the infections, the parasitaemia of mice infected with single infections of *Pv*DS were significantly lower than those of mice infected with *Pv*DS+*Pc*AJ mixed-infections (two-way RM ANOVA measured mixed effects model, F=218.8, DFn=1, DFd=3, P=0.0007). The erythrocyte density of mice infected with *Pv*DS+*Pc*AJ mixed-infections was significantly lower than those infected with single infections of *Pv*DS (two-way RM ANOVA measured mixed effects model, F=123, DFn=1, DFd=3, P=0.0016). Detailed statistical values relating to significance are given in Supporting Table 1. Experiments were repeated twice, data is from one representative experiment.

### Co-infection of *P. yoelii* with either *P. chabaudi* or *P. vinckei* results in reduced parasite density of *P. yoelii*, and protracted peak parasitaemia of *P. chabaudi* and *P. vinckei*

To understand how mixed parasite species infection influences the fitness of the species involved, we measured the parasite density (numbers of parasites per ml of mouse blood) through time of individual species in single and mixed infections. The relative proportions of each species within mixed infections were measured at 24-hour intervals by species specific qPCR.

In mixed infections composed of *Py*CU and *Pv*DS*, Pv*DS dominated the infection from days 4 to 10 PI (Fig. 3A), at which point *Py*CU became dominant. There was an increase in the proportion of *Py*CU in the infection from day 8 (7%) until host mortality at day 12 PI (50%). Analysis of species-specific parasite density in this co-infection revealed that *Py*CU was suppressed throughout the infection, while the growth of *Pv*DS was enhanced (Fig. 3D, E). This enhancement occurred during the latter stages of the infection (days 8-12), the time point at which *Pv*DS is cleared during single infections. This suggests that the presence of *Py*CU, whose growth in a co-infection does not differ significantly from that observed in a single species infection, facilitates the persistence of *Pv*DS for an extended period after which it would normally be cleared. This inability to clear *Pv*DS, combined with the standard increase in *Py*CU parasitaemia, leads to hyper-parasitaemia with severe anaemia in co-infected mice, and results in host death.

**Fig. 3.**
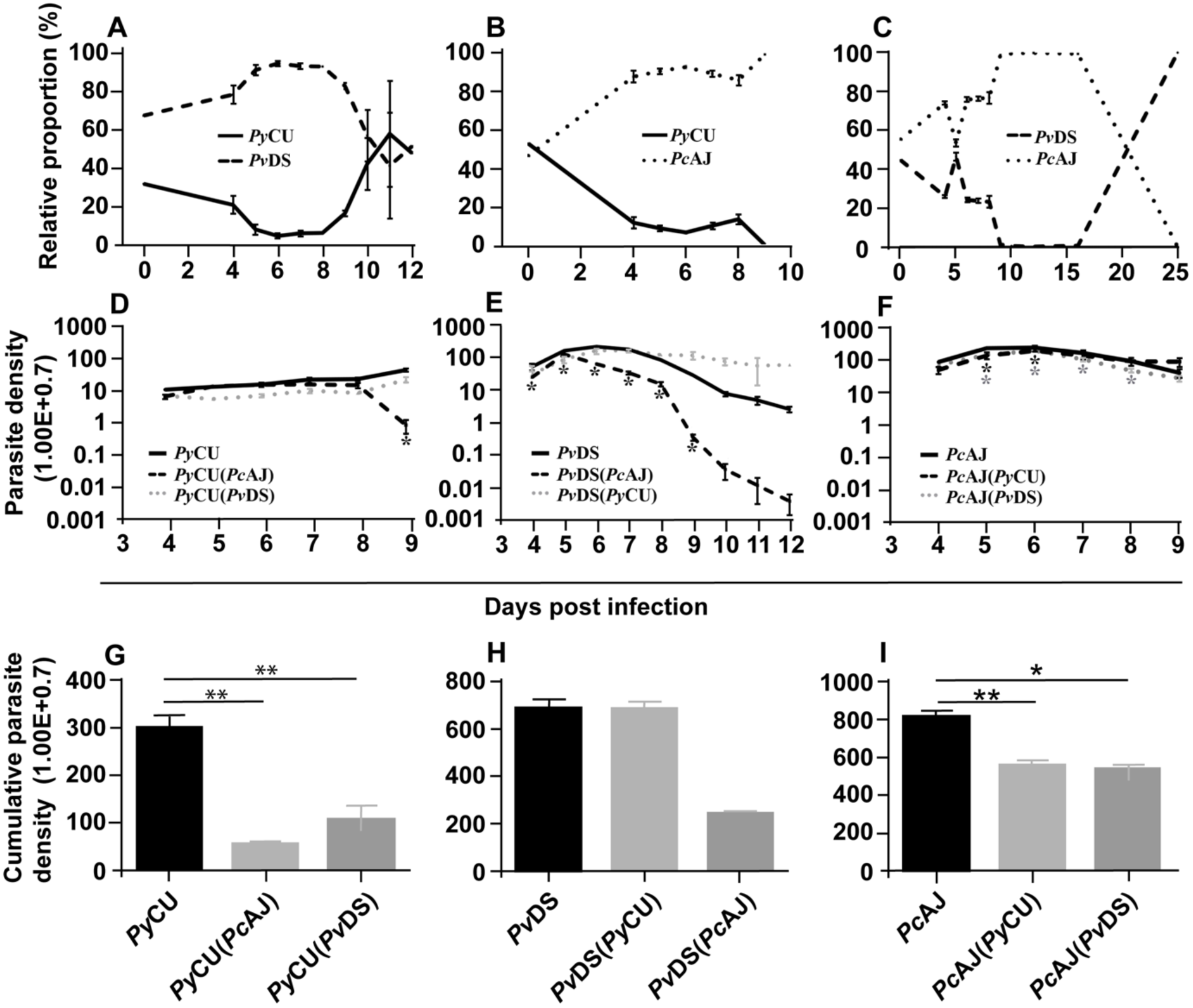
The relative proportions of *Plasmodium yoelii* CU, *Plasmodium vinckei* DS, and *Plasmodium chabaudi* AJ in mixed infections (panels A-C), the parasite density of each species in single and mixed infections (panels D-F), and cumulative parasite density of each species in either single or mixed infections (panels G-I). The relative proportion of each species in combination with each other (panels A-C) was measured by qPCR quantification using primers specific to a region of the *msp1* gene of each species. Copy numbers of parasite *msp1* were quantified with reference to a standard curve generated from known numbers of plasmids containing the same gene sequences. The average copy numbers per iRBC were generated by copy numbers and parasite densities of each single species infections on day 6 PI. Data points indicate the mean value for 3 to 5 mice in each experimental group and error bars indicate the standard error of the mean (SEM). The parasite densities (number of blood stage parasites per mL blood) are shown in panels D-F. Panel D shows the parasite densities of *Py*CU single infection or in mixed infection with *Pv*DS or *Pc*AJ. Parasite densities of PvDS and PcAJ are given in panels E and F, respectively. An asterisk represents significant differences (P < 0.01) in parasite density in single infections compared with mixed-infections. Cumulative parasite densities are shown in panels G-I. The cumulative parasite density of *Py*CU in single infections were significantly higher than *Py*CU in mixed-infections with *Py*AJ and *Pv*DS (panel G). Similarly, the cumulative parasite density of *Pc*AJ in single infections was significantly higher than in mixed infections with *Py*CU or *Pv*DS (panel I). Detailed statistical values relating to significance are given in Supporting Table 1. Experiments were repeated twice, data is from one representative experiment.

Enhancement of the parasite density of the normocyte-restricted parasite species was also observed in the latter stages of mixed species infections composed of *Py*CU and *Pc*AJ. In this case, *Pc*AJ dominates *Py*CU throughout the infection, with complete exclusion of the latter species observed at the end of the co-infection (Fig. 3B). Mice died at days 9 and 10 pi, at which point the parasite density of *Py*CU was significantly suppressed compared to single infections (Fig. 3D). In contrast, there was little difference in the parasite density of *Pc*AJ in a mixed infection with *Py*CU compared to a single infection during the first eight days of the co-infection. However, as seen in the *Pv*DS+*Py*CU infection, the usual reduction in parasitaemia observed at day 8 in *Pc*AJ single infections was not observed in co-infections with *Py*CU (Fig. 3F), suggesting again that presence of *Py*CU in a co-infection impairs the ability of the host to control the growth of the normocyte-invading parasite species.

There was a dramatic reduction in parasite density for *Py*CU in the mixed infections with both *Pc*AJ and *Pv*DS (Fig. 3G). In the mixed infections involving *Pc*AJ and *Pv*DS, both of which preferentially invade normocytes, there was a dramatic reduction in parasite density for *Pv*DS throughout the co-infection compared to single infection (Fig. 3H). There was also a slight reduction in parasite density of *Pc*AJ (Fig. 3I), especially during the latter stages of the infection. *Pc*AJ is the more virulent of the two species and dominates the co-infection between days 5 and 15, when no *Pv*DS could be detected by qPCR. However, *Pv*DS resurges at day 15, and competitively excludes *Pc*AJ by day 25.

### Pre-exposure of mice to *P. yoelii* does not enhance the virulence of *P. vinckei* infection

As the presence of *P. yoelii* in mixed infections with either *P. chabaudi* or *P. vinckei* results in an inability to clear the latter two species, resulting in host death, we wondered whether an immune response specific to *P. yoelii* could adversely affect the establishment of an effective immune response against *P. chabaudi* or *P. vinckei*. To test this, we pre-immunised mice with *Py*CU parasites by exposure to the parasite for eight days followed by clearance with the anti-schizontal drug mefloquine (MF). Two weeks later, when all MF had cleared from the host, mice were challenged with *Pv*DS. In contrast to the patterns observed in *Py*Cu+*Pv*DS co-infections, there was no evidence of increased virulence of *Pv*DS infections in mice pre-exposed to *Py*CU (Fig. 4A), with no significant enhancement of parasite density (Fig. 4B), anaemia (Fig. 4C), or weight loss (Fig. 4D) occurring at any stage during the infection.

**Fig. 4.**
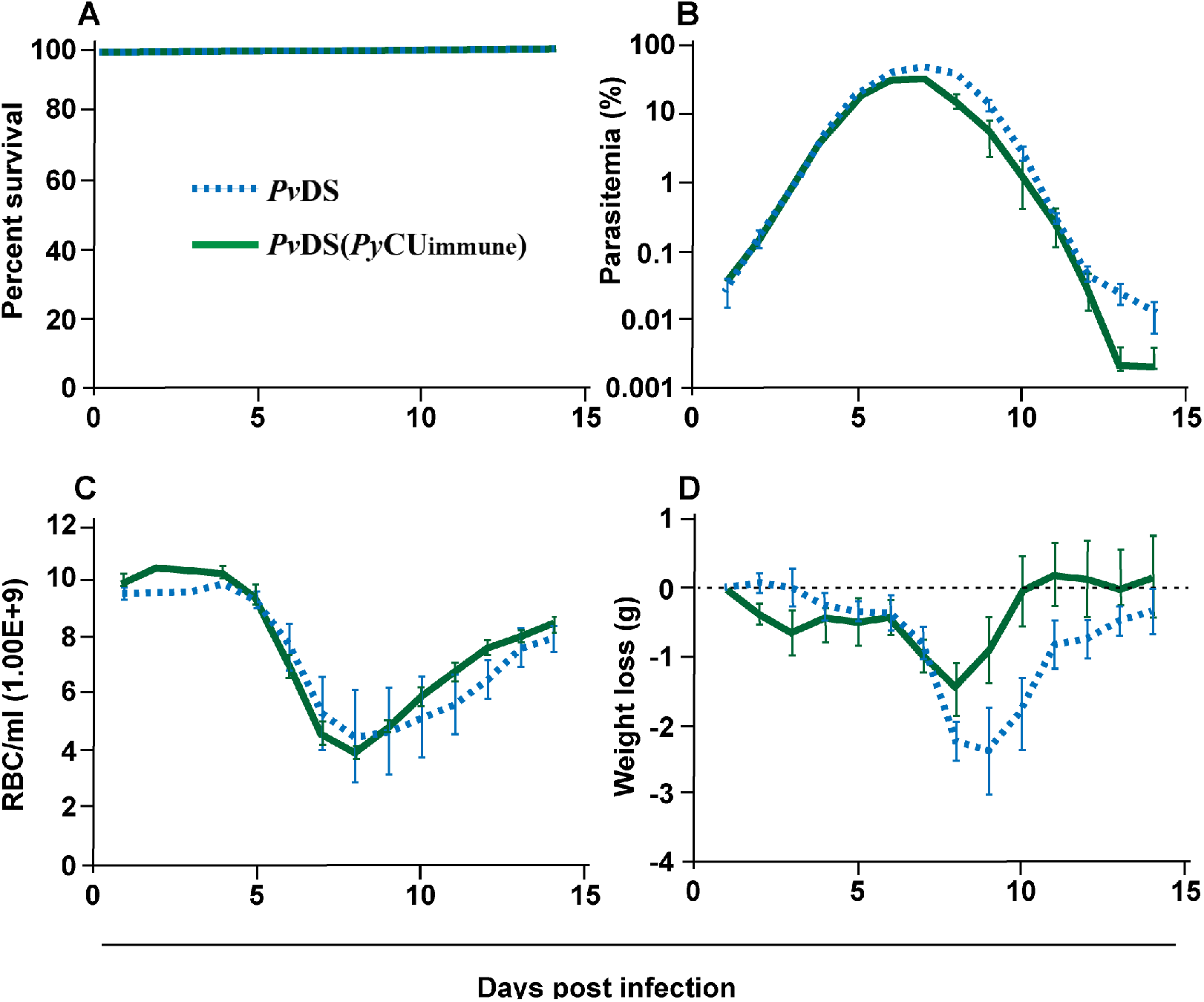
Percentage survival (panel A), parasitaemia (panel B), erythrocyte density (panel C), and weight loss (panel D) of mice infected with *Plasmodium yoelii* CU (*Py*CU) and *Plasmodium vinckei* DS (*Pv*DS) in single and mixed infections. Mice in the mixed-infection group were inoculated intravenously (IV) with 1×10^6^ *Py*CU infected red blood cells (iRBCs) seven days prior to IV inoculation with 1×10^6^ *Pv*DS iRBCs. Data points indicate the mean value for 4 to 5 mice in each experimental group and error bars indicate the standard error of the mean (SEM). Asterisks indicate statistically significant differences between mice with mixed-species infections compared with both single infections (Multiple t-tests, with the assumption that all rows are sampled from populations with the same scatter and corrected for multiple comparisons using the Holm-Sidak method). Mice infected with mixed-infections lost significantly more weight during the latter stages of the infection than mice infected with single species infections (days 21-23 PI, panel D). The relative proportion of each species in mixed infections (panel E) was measured by qPCR quantification of the *msp1* gene. The average copy number per iRBC was generated with reference to copy numbers and parasite densities of each single species infections on day 6 PI. The parasite densities of *Py*CU and *Pv*DS in single or mixed infections are shown in panels F and G. A Mann Whitney test shows that the cumulative parasite density of *Pv*DS in single infections is significantly higher than that of *Pv*DS in a mixed infection with *Py*CU (panel I), whereas that of *Py*CU is unaffected when in a mixed infection with *Pv*DS (panel H). Detailed statistical values relating to significance are given in Supporting Table 1.

### The increased virulence of mixed species infections of *P. yoelii* and *P. vinckei* is abrogated when *P. vinckei* is added to an established *P. yoelii* infection

When both *Py*Cu and *Pv*DS are inoculated into mice contemporaneously, the resulting co-infection is consistently lethal, in contrast to the zero-mortality associated with the constituent single species infections. This lethality results from the inability of mice to clear the *Pv*DS parasites from the circulation following peak parasitaemia. To determine if increased virulence is dependent on the timing of the introduction of the co-infecting species, we first inoculated mice with *Py*CU and introduced *Pv*DS seven days later. The co-infection caused 25% mortality, compared to no mortality in single species infections (Fig. 5A). In this case, the co-infection parasitaemia did not differ significantly from that of a single infection of *Py*CU for most of the infection duration, except for the last two days of sampling (days 22 and 23), when the parasitaemia was higher in the co-infection (Fig. 5B). This increased parasitaemia towards the latter stages of the infection did not result in lower erythrocyte density (Fig. 5C); however, it did cause significantly greater weight loss in co-infected animals during this period (Fig. 5D).

**Fig. 5.**
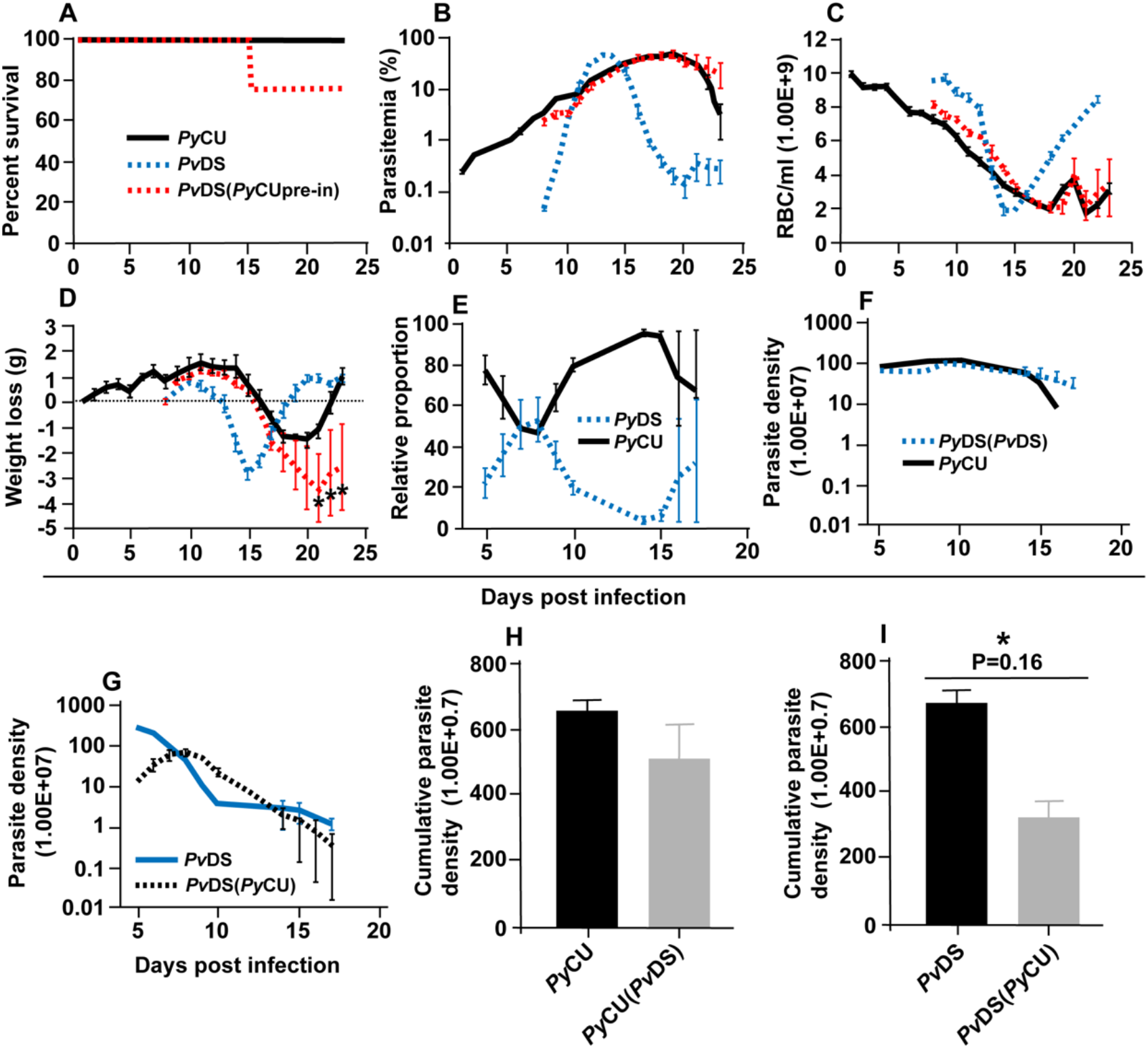
Percentage survival (panel A), parasitaemia (panel B), erythrocyte density (panel C), and weight loss (panel D) of mice infected with *Plasmodium vinckei* DS (*Pv*DS). Mice in the *Pv*DS (*Py*CU immune) group were inoculated intravenously (IV) with 1×10^6^ infected red blood cells (iRBCs) of *Py*CU, treated 8 days later with mefloquine for 5 days, and then 15 days later intravenously challenged with 1×10^6^ *Pv*DS iRBCs. Data points indicate the mean value for 5 mice in each experimental group and error bars indicate the standard error of the mean (SEM). Detailed statistical values relating to significance are given in Supporting Table 1.

Measuring the relative proportions and parasite densities of the constitute species in the co-infection and comparing them to single infections revealed that *Py*CU dominates the infection over *Pv*DS, excepting days 7 and 8 (Fig. 5E). In contrast to the situation observed with the simultaneous inoculation of the two species, there was no significant reduction in the parasite density of *Py*CU, but there was a reduction in the parasite density of *Pv*DS when *Py*Cu was inoculated one week prior to *Pv*DS (Fig. 5F-I).

### The consequences of mixed species infections in the mosquito vector

We additionally sought to describe the impact of mixed species infections on transmission to mosquitoes. Specifically, we fed *Anopheles stephensi* mosquitoes on mice with single or mixed species infections and measured: i) the proportion subsequently infected, and ii) the severity of this infection (number of oocysts); iii) the longevity of infected mosquitoes; and iv) the number of larvae they produced following a blood meal. Finally, we compared the transmission success, defined as the average number of oocysts produced per blood fed mosquito, of each parasite species in mixed or single infections.

### Mixed species infections do not result in significantly different infection parameters in mosquitoes

To determine whether mixed species infections result in altered mosquito infectivity rates and infection loads compared to single species infections, mosquitoes were fed on anaesthetized mice infected with single, or mixed infections of *Pc*CU, *Pv*DS, and *Pc*AJ. As these species differ in the timing of their gametocyte production, with *Py*CU at its most infectious to mosquitoes on day 3 PI and *Pc*AJ and *Pv*DS more infectious on day 5 pi, we conducted mosquito feeds on both these days. *Py*Cu was the most infective single species on day 3 and day 5, followed by *Pv*DS and finally *Pc*AJ (Fig. 6A-C). Mixed species infections did not result in higher proportions of mosquitoes being infected than the most infective constituent species in a single infection (Fig. 6A-C). Similarly, in co-infections containing the highly infectious *Py*CU species, the oocyst burdens of mixed species infections were not significantly different from that of *Py*CU in mosquitoes fed on mixed infections on either day 3 or day 5 PI (Fig. 6D+E). Mixed species infections of *Pc*AJ and *Pv*DS resulted in significantly lower oocyst burdens than the most infectious constituent single species (*Pv*DS), but only in mosquitoes fed on day 5 of the infection (Fig. 6F).

**Fig. 6.**
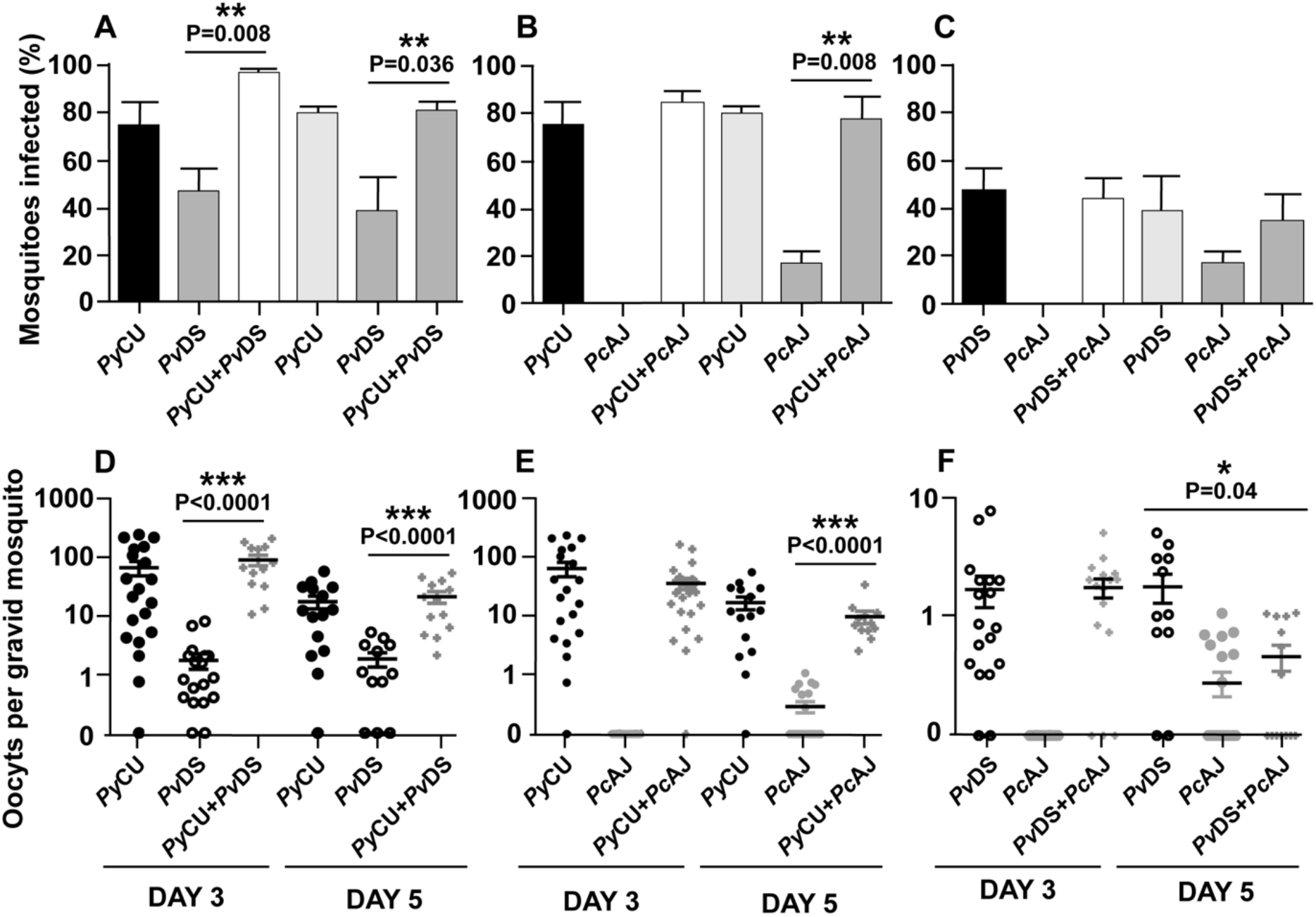
Survival (panels A-D) and larvae per mosquito (panel E) of mosquitoes infected with single and mixed species infections. Mosquito survival curves, infected with either single species or mixed infection of *Plasmodium yoelii* CU, *Plasmodium vinckei* DS, and *Plasmodium chabaudi* AJ are shown in panels A-C. Longevity of infected mosquitoes was observed until day 60 after the blood meal and the number of dead mosquitoes were recorded every 5 days. Boxplots indicate median survival and first and third quartiles, and whiskers are the same quartiles ± (1.5×interquartile range) (panel D). The numbers of larvae from infected mosquitoes with single or mixed infections were recorded and no significant difference was observed. Detailed statistical values relating to significance are given in Supporting Table 1.

### Co-infections of malaria parasite species do not adversely affect mosquito longevity or capacity to produce larvae

To ascertain whether mixed species infections of mosquitoes were more virulent than single species infections, we measured longevity and larvae production in mosquitoes fed on single or mixed infections. In accordance with the observation that mixed species infections did not result in higher burdens of infection, mosquitoes infected with two parasite species in a co-infection did not display reduced longevity (Fig. 7A-C), median survival time (Fig. 7D), or reduced fitness (measured as the number of larvae produced per blood-fed mosquito) (Fig. 7E).

**Fig. 7.**
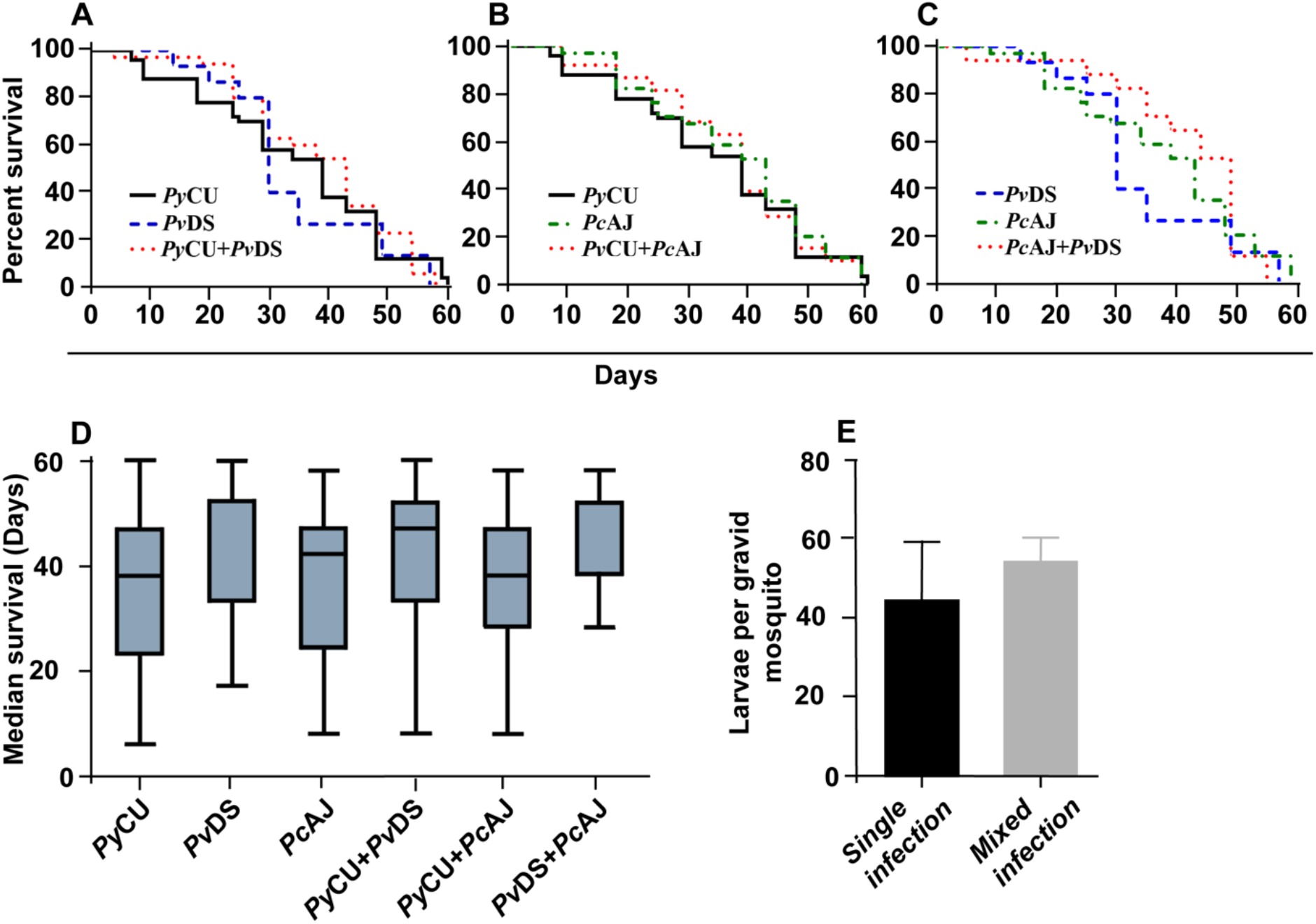
Transmission capacity of *Plasmodium yoelii* CU (*Py*CU), *Plasmodium vinckei* DS (*Pv*DS), and *Plasmodium chabaudi* AJ (*Pc*AJ) in single or mixed infections. The mean percentage of mosquitoes infected with oocysts were calculated following feeding on *Py*CU, *Pv*DS, and *Pc*AJ either in single or mixed infections in mice on days 3 and 5 PI (panels A-C). Only gravid mosquitoes were considered blood-fed and included in the analysis. Statistical analysis was performed using Mann Whitney tests. Detailed statistical values relating to significance are given in Supporting Table 1.

### Mixed species infections can affect the transmission capacity of the constituent species

To determine whether mixed infections can affect the transmission capacity of constituent species, the relative proportion of each species in mixed infections in mosquitoes was measured using qPCR on DNA extracted from mosquitoes with known oocyst numbers and compared to the numbers produced in single infections.

The numbers of oocysts produced by the highly infectious *Py*CU did not differ significantly between mosquitoes fed on single and mixed species infections (Fig. 8A). *Pv*DS produced fewer oocysts in mixed infections with *Pc*AJ and *Py*CU than in single infections, although the effect was only statistically significant on day 5 in a mixed infection with *Py*CU (Fig. 8B). *Pc*AJ also suffered a reduced transmission capacity in mixed infections, with significant reductions in oocyst numbers measured in mosquitoes fed on mixed infections with either *Pv*DS or *Py*CU on day 5, compared to those fed on single infections (Fig. 8C). The transmission of *Pc*AJ to mosquitoes was completely blocked in mixed infections containing *Py*CU (Fig. 8C).

**Fig. 8.**
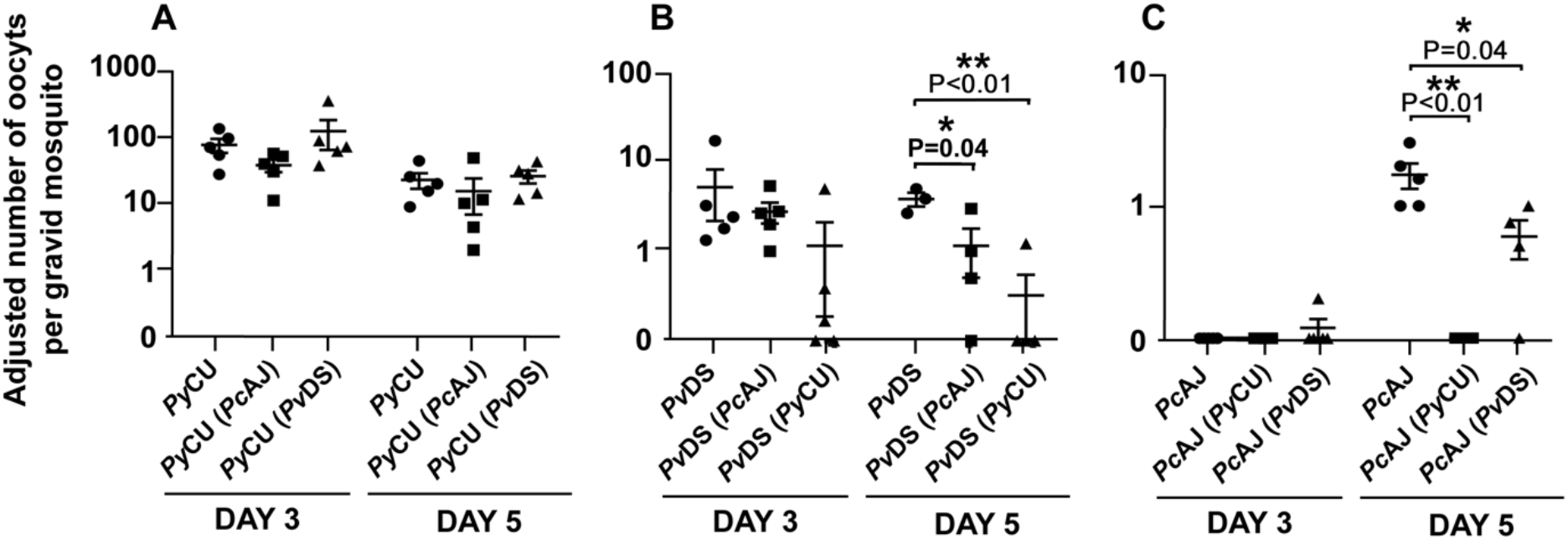
Adjusted mean number of oocysts of *Plasmodium yoelii* CU (*Py*CU), *Plasmodium vinckei* DS (*Pv*DS), and *Plasmodium chabaudi* AJ (*Pc*AJ) in single or mixed species infections per gravid mosquito. Data points represent the mean oocyst burden of mosquitoes fed on individual mice (n = 5 per group). There was no statistical difference between *Py*CU in single and in mixed infections with either *Pc*AJ or *Pv*DS (panel A), *Pv*DS was suppressed when mixed with *Py*CU on day 5 PI (panel B) and *Pc*AJ was suppressed in mixed infections with *Py*CU and *Pv*DS on day 5 PI (panel C). Detailed statistical values relating to significance are given in Supporting Table 1.

## Discussion

The major malaria parasite species that naturally infect man have overlapping ranges in many tropical regions, the greatest exception being caused the apparent absence of *P. vivax* from large parts of west and central and central Africa, where *P. ovale*, *P. malariae* and *P. falciparum* are at their most abundant. It is often difficult to interpret data regarding the consequences of the mixed-species malaria infections that occur naturally in man, due to the multitude of confounding factors that exist in nature.

Our results indicate that the interactions between malaria parasites co-infecting the same host can have dramatic consequences for the severity of the disease they cause. We found that when two parasite species, *P. yoelii* and *P. vinckei*, which on their own cause mild and transient disease concurrently infect the same host, the disease outcome is radically altered resulting in 100% host mortality within 15 days. This same outcome of 100% host mortality was observed in co-infections consisting of *P. yoelii* and the more virulent (but rarely lethal) *P. chabaudi*.

There are precedents for this result; Bafort (1971) suggested that mixed species might increase the virulence of infections [36]. Richie (1985) reported that patent *P. chabaudi* infections increased their parasitaemia and duration when mixed with *P. yoelii* [9], and McGhee (1964) also described a higher peak for one of the species in mixed infections [37]. More recently, Ramiro *et al* (2016) described increased virulence in mixed infections of *P. chabaudi* and *P. yoelii*, which they attributed to an increase in reticulocytaemia leading to enhancement of *P. yoelii* (which is reticulocyte restricted) in the mixed infections. Our results, however, are in agreement with those of Richie (1988), and suggest that it is the normocyte-restricted parasite that is enhanced in mixed infections with *P. yoelii*, a result we observed both in the case of two strains of *P. chabaudi* and one of *P. vinckei*. We contend, therefore, that increasing reticulocytaemia does not explain the increase in virulence of mixed strain infections. This is supported by the fact that we also observed increased virulence (in terms of persistence of infection and decreased red blood cell density) in mixed-species infections composed of *P. chabaudi* and *P. vinckei*, both of which predominantly infect normocytes.

We found that the time at which the constituent species of a mixed-species infection were introduced to the host had a significant impact on disease outcome. Most studies on mixed malaria parasite species infections in mice introduce the constituent species contemporaneously [27, 28]. For example, the parasite species combination *P. yoelii* and *P. vinckei* causes 100% mortality when introduced to mice contemporaneously. We found that when *P. yoelii* was inoculated seven days prior to the inoculation of *P. vinckei*, virulence was much reduced, although the mixed infection still caused significantly more pathology and mortality than the constituent species in single infections. Similarly, when inoculated into mice contemporaneously with *P. vinckei*, *P. yoelii* suffered a reduction in cumulative parasite density throughout the infection (a proxy measurement of parasite fitness), whereas *P. vinckei* was unaffected. When *P. yoelii* was introduced to mice a week before *P. vinckei*, the opposite trend was observed, with little reduction in the cumulative parasite density of *P. yoelii*, but a significant reduction in that of *P. vinckei*. We show, therefore, that it is not just the phenotypes of the constituent species of mixed infections that affect pathology and parasite fitness in combination, but also the time at which each species infects the host.

It is possible that interactions between malaria parasite species in mixed infections may be modulated through the host immune response. Molineaux (1980) [38] suggested that in mixed infections of *P. falciparum* and *P. malariae*, the immune response stimulated by rising *P. falciparum* parasitaemias can inhibit *P. malariae*, but that *P. falciparum* survives longer due to its more rapid growth rate. This immune-mediated antagonism [39] agrees with observations suggesting that *P. falciparum* could reduce the prevalence of *P. malariae* [40].

Compelling evidence suggests cross-protection between malaria parasite species due to species-transcending immunity [41, 42]. We wondered whether the lack of ability to control the *P. vinckei* parasitaemia towards the end of the mixed infection of *P. vinckei* and *P. yoelii* may be due to the phenomenon of “original antigenic sin” [43] rendering the acquisition of antibodies specific to *P. vinckei* sub-optimal due to the larger quantity of *P. yoelii* antigen present in the early stages of the infection. To test this, we immunised mice through exposure to and subsequent cure of a *P. yoelii* infection, and then challenged with *P. vinckei*. Contrary to the expectations of the original antigenic sin hypothesis, we observed no effect on the severity of the *P. vinckei* infection in *P. yoelii*-exposed compared to non-exposed mice.

Infection with malaria parasites is known to detrimentally affect the fitness of the infected mosquito [44]. Mixed species malaria parasite infections are common in nature [4, 45], and as mixed species parasite infections caused dramatically different disease outcomes in mice, we investigated whether mosquito fitness was also affected. We measured longevity and progeny production in groups of mosquitoes fed on single and mixed species infections. In contrast to the significant alterations in pathogenicity observed in mice, there appeared to be no fitness differences between mosquitoes carrying single or mixed species infections. Linked to this, we did not observe significantly increased oocyst numbers in mosquitoes infected with mixed species, when compared to the highest-oocyst producing single constituent species, suggesting there was no significant alteration in transmission-stage investment by the species in mixed stage infections [31].

We found that the transmissibility of *P. vinckei* and *P. chabaudi* to mosquitoes was reduced in the presence of *P. yoelii* in co-infections compared to single infections. This was reflected in the lower number of oocysts of these two species in mosquitoes that had fed on mixed species infections also containing *P. yoelii* compared to those that had fed on single infections. Most significantly, transmission of *P. chabaudi* to mosquitoes was blocked completely by the presence of *P. yoelii* on day 5 of the infection. There were no reductions, however, in the numbers of *P. yoelii* oocysts. Of the three species, *P. yoelii* produces significantly higher oocyst burdens in mosquitoes than either *P. chabaudi* or *P. vinckei*, a phenomenon linked to the former species having much higher gametocyte production during the early stages of infection. One possible mechanism that may account for this observation involves gamete incompatibility; we propose that given the fact that in a mixed species infection containing *P. yoelii*, there will be significantly more *P. yoelii* microgametes than of the other species. If *P. yoelii* microgametes can recognise and attempt to fertilise the macrogametes of the second species, then a large proportion of these macrogametes will be rendered non-productive (assuming hybrids are non-viable) [46]. Consistent with this theory is the fact that both *P. chabaudi* and *P. vinckei* also produce fewer oocysts when in mixed infection with each other, but much less so than when mixed with *P. yoelii*, reflecting, perhaps, the more even numbers of gametocytes produced by these two species.

As the world moves towards reducing the malaria burden, the importance of mixed-species malaria infections will rise. Diagnostic techniques with improved sensitivities are revealing a greater prevalence of *P. ovale* and *P. malariae* in *P. falciparum* endemic areas than previously thought [7]. Intervention strategies such as anti-vector programs, and the development and employment of new drugs and vaccine will often be more effective against one species of parasite than they will against others [47, 48]. In regions where *P. falciparum* prevalence is decreasing, the prevalence of non-falciparum malaria parasite species often becomes more apparent, highlighting the importance of mixed-species infections. There is a need, therefore, to better understand how the interactions between malaria parasites species infecting the same host can impact disease progression and parasite fitness.

The experiments described here show that the disease outcomes of mixed versus single species infections can differ, and are influenced by the phenotypic characteristics of the constituent species and the order in which they infect the host.

## Acknowledgments

RC is supported by Japanese Society for the Promotion of Science (JSPS), Japan Grant-in-Aid for Scientific Research No. 19K07526. JT is supported by the Jiangsu Provincial Department of Science and Technology Grant (No. BM2018020) and The Jiangsu Provincial Project of Invigorating Health. Care through Science, Technology and Education

## Author Contributions Statement

Designed, performed and analysed the experiments; RC and JT. Wrote the manuscript; RC, JT, TJT and JC.

